# Antimicrobial resistance gene lack in tick-borne pathogenic bacteria

**DOI:** 10.1101/2022.11.28.518145

**Authors:** Márton Papp, Adrienn Gréta Tóth, Gábor Valcz, László Makrai, Sára Ágnes Nagy, Róbert Farkas, Norbert Solymosi

## Abstract

Tick-borne infections, including those of bacterial origin, are significant public health issues. Antimicrobial resistance (AMR), which is one of the most pressing health challenges of our time, is driven by specific genetic determinants, primarily by the antimicrobial resistance genes (ARGs) of bacteria. In our work, we investigated the occurrence of ARGs in the genomes of tick-borne bacterial species that can cause human infections. For this purpose, we processed short/long reads of 1550 bacterial isolates of the genera *Anaplasma* (n=20), *Bartonella* (n=131), *Borrelia* (n=311), *Coxiella* (n=73), *Ehrlichia* (n=13), *Francisella* (n=959) and *Rickettsia* (n=43) generated by second/third generation sequencing that have been freely accessible at the NCBI SRA repository. From *Francisella tularensis*, 98.9% of the samples contained the FTU-1 gene, and 16.3% contained additional ARGs. Only 2.2% of isolates from other genera (*Bartonella*: 2, *Coxiella*: 8, *Ehrlichia*: 1, *Rickettsia*: 2) contained any ARG. We found that the odds of ARG occurrence in *Coxiella* samples were significantly higher in isolates related to farm animals than from other sources. Our results describe a lack in ARGs in these bacteria and suggest that antibiotic susceptibility testing might be considered before the treatment of tick-borne infections in farm animals.

## Introduction

While antimicrobial resistance (AMR) is often referred to as one of the main challenges of the 21st century^1^, climate change, urbanization, and globalization may also bring unprecedented changes in public health, namely by affecting the population size and geographical distribution of various disease vectors^2^. Several studies prove the widespread dissemination of antimicrobial resistance genes (ARGs) in various environmental or alimentary sample types^3–7^. Whilst AMR and its underlying causes, ARGs are distributed worldwide and thus may affect human and animal populations globally, vector-borne diseases are also not uncommon. Indeed, around 80% of the human population of the Earth is estimated to be at risk of one or more vector-borne diseases (VBDs)^8^. Since many of these VBDs are bacterial, treating a set of vector-borne infections relies upon the efficacy of antibiotics. Despite the potential and relatively dynamic changes expected in the spread of both AMR and VBDs in the coming decades, the associations of these global health issues are less studied.

Besides such significant arthropod disease vectors as mosquitoes, sandflies, blackflies, fleas, lice, tsetse flies, or triatome bugs, soft and hard ticks may also serve as vectors for pathogenic microorganisms of human and veterinary medical significance^8,9^. Considering that the spectrum of tick-borne bacterial diseases is relatively wide, the assessment of the antimicrobial resistance status of tick-borne pathogens, such as the members of the genera *Anaplasma, Bartonella, Borrelia, Coxiella, Ehrlichia, Francisella* and *Rickettsia* is an anticipatory public health step^10–16^.

Our study aimed to obtain detailed knowledge of the ARG spectrum of the bacteria that cause the primary tick-borne infections. Therefore, whole-genome sequencing data of bacterial isolates belonging to the genera of *Anaplasma, Bartonella, Borrelia, Coxiella, Ehrlichia, Francisella, Rickettsia* were processed using a uniform bioinformatics pipeline.

## Materials and Methods

### Samples

The available samples and corresponding sequencing data for each genus were retrieved and downloaded from the SRA database on the following days: *Anaplasma* Aug 16, *Bartonella* Aug 19, *Borrelia* Aug 21, *Coxiella* Aug 27, *Ehrlichia* Aug 12, *Francisella* Aug 12, *Rickettsia* Aug 12. Of the downloaded samples, the number of WGS samples per genus was as follows: *Anaplasma*: 40, *Bartonella*: 251, *Borrelia*: 707, *Coxiella*: 155, *Ehrlichia*: 16, *Francisella*: 1088, *Rickettsia*: 128.

For further analysis, the raw sequencing data from Illumina, Nanopore or PacBio platforms were considered only. Moreover, the analysis was restricted to samples from species of potential tick-borne spread and associated human diseases. De novo assembly was performed on the sequencing data of these samples, followed by the prediction of the ARG content of the generated contigs. Only samples with an available representative genome at the NCBI database were included with the condition that sequencing coverage reached at least 95% of the genome. After the selection steps, a total of 1550 samples (*Anaplasma*: 20, *Bartonella*: 131, *Borrelia*: 311, *Coxiella*: 73, *Ehrlichia*: 13, *Francisella*: 959, *Rickettsia*: 43) were left for analysis (BioSample IDs of the samples at each genus considered for the analysis can be found in Supplementary File 1 and 8).

### Bioinformatic analysis

For Illumina sequenced data, the quality-based filtering and trimming of the raw short reads was performed by TrimGalore (v.0.6.6, https://github.com/FelixKrueger/TrimGalore), setting 20 as the quality threshold. Only reads longer than 50 bp were retained. Nanopore sequenced reads were adapter trimmed and quality-based filtered by Porechop (v0.2.4, https://github.com/rrwick/Porechop) and Nanofilt (v2.6.0)^17^, respectively. The long reads from the PacBio sequencing platform were prepared for assembly using the pbclip (https://github.com/fenderglass/pbclip) tool.

The Illumina reads were assembled to contigs by MEGAHIT (v1.2.9)^18^ using default settings. The assembly of the long read sequencing data was performed with Flye (v2.9-b1779)^19^, and 3x round polishing was applied with Racon (v1.4.3)^20^ on the resulting contigs. In the case of hybrid sequencing, the sequenced reads were assembled by MaSuRCA (v4.0.9)^21^.

All possible open reading frames (ORFs) were predicted by Prodigal (v2.6.3)^22^ on each contig. The protein-translated ORFs were searched for ARG sequences against the Comprehensive Antibiotic Resistance Database (CARD, v.3.2.3)^23,24^ by the Resistance Gene Identifier (RGI, v5.2.0), running with Diamond^25^ as the aligner. To capture ARGs with high-confidence rates for the following steps of the analysis, ORFs classified as perfect or strict matches were further filtered by 90% identity and 90% coverage thresholds.

ARGs predicted by RGI were further screened for potential associated mobile genetic elements to asses their potential for participation in Horizontal Gene Transfer (HGT) events. Contigs, throughout which ARGs were identified, were screened for the probability of plasmid origin by PlasFlow (v.1.1)^26^. The prediction of integrative mobile genetic elements (iMGE) on the contigs harboring ARGs were assessed by the MobileElementFinder (v1.0.3) and its database (MGEdb, v1.0.2)^27^. An ARG was considered to be associated with an iMGE if it was within the distance of the median for the longest composite transposon stored for each bacterial species in the MGEdb database (distance threshold: 10098 bp). Bacteriophage sequences were predicted with VirSorter2 (v2.2.3)^28^ and were filtered to dsDNAphage and ssDNA sequences only. ARGs within phage sequences at their whole length were considered to be associated with bacteriophages.

To mitigate the effect of the database and tool used for the ARG prediction on our results, we have further analysed them with two additional pipelines. AMRFinderPlus (v.3.10.36)^29^ is an alignment-based tool similar to RGI. We used AMRFinderPlus with a database (National Database of Antibiotic Resistant Organisms, NDARO, version 2022-05-26.1) without the --plus option to retain as much comparability as possible between its results and those from RGI and CARD. Similarly, the determined genes were filtered by 90% identity and 90% coverage thresholds. KmerResistance^30^ aligns raw reads to a user-defined database with the aid of the KMA aligner^31^ while also handling potential contamination. We have searched only the Illumina sequencing data from the samples considered in our analysis (for the samples included in the KmerResistance analysis, see Supplementary File 1) with KmerResistance (v2.0) against the CARD (v3.2.3) and NDARO (v2022-05-26.1) databases. Therefore, the number of samples analysed with KmerResistance as well was 1492. From the CARD database, only genes under the protein homolog models were used for the KmerResistance analysis. Results were filtered by 90% template coverage and 90% query identity thresholds. As the AMRFinderPlus analysis was run without the --plus option, virulence genes and genes confering resistance to metal- and biocide compounds only were filtered from the KmerResistance results when run against the NDARO database.

The association of the origin of the samples and the ARG positivity was analysed based on the metadata available for each biosample at the SRA database. The origin of samples was determined by the sample_origin and sample_host metadata columns. Based on this metadata information, the samples were manually sorted into the following categories: companion animal, environment, farm animal, human, vector, wild animal and NA when no information was available. Also, an NA category was used in the case of indicated hosts of frequently used laboratory animal species, as in those cases, we couldn’t determine if the host and the sample were experimentally manipulated. Additionally, if any other metadata indicated the laboratory origin of the strain, the NA category was used for the sample origin.

## Results

The ARG content of available WGS samples of human-pathogenic bacteria with possible tick-borne origin from the SRA database was analysed. After selecting samples with raw data of sufficient quality (for details, see Materials and Methods), 1550 samples remained for the ARG analysis. Samples represent species from 7 bacterial genera (*Anaplasma, Bartonella, Borrelia, Coxiella, Ehrlichia, Francisella* and *Rickettsia*), originating from at least 33 different countries. Even though most samples were supplemented with metadata of their country of origin, there were still 142 samples with this information missing. Countries with at least one sample included in our analysis are shown in Figure 1. Predicted ARG content was analysed with RGI and CARD, a robust pipeline often employed to survey antimicrobial resistance genes in different environments^3–7^. However, as resistance gene presence was rather scarce in the samples analysed in this study, we have included two additional methods for resistance gene prediction to augment our results. Even though we utilized these additional methodologies, we base our findings on the widely used and accepted RGI pipeline. Further information about the samples can be found in Supplementary File 1.

**Figure 1.**
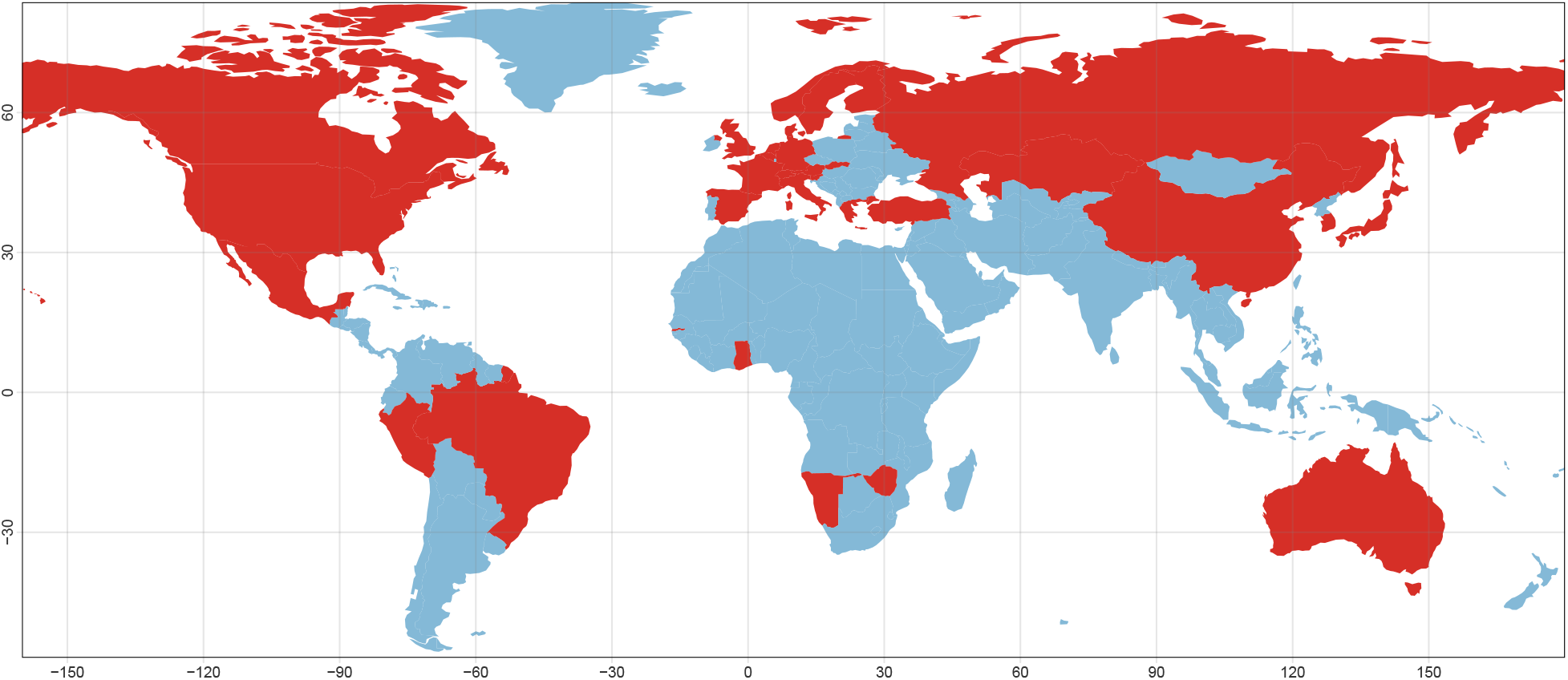
World map indicating the countries with at least one sample included in the analysis (red area). The countries with samples are the following: Australia, Austria, Belgium, Brazil, Canada, China, Denmark, Finland, France, Gambia, Germany, Ghana, Greece, Italy, Japan, Kazakhstan, Liechtenstein, Mexico, Namibia, Netherlands, Norway, Peru, Russia, Slovakia, Slovenia, South Korea, Spain, Sweden, Switzerland, Turkey, UK, USA, Zimbabwe. There were 142 samples without a designation on their country of origin.

### Resistance genes predicted by RGI and CARD

Even though we could include a large number of samples in our analysis, a surprisingly low number of resistance genes were predicted from the ORFs. Furthermore, the ARGs were distributed among only a handful of samples. The percentage of ARG positive samples was 0% for the *Anaplasma* and *Borrelia* genera, 1.5% for the *Bartonella* genus, 11.0% for the *Coxiella* genus, 7.7% for the *Ehrlichia* genus and 4.7% for the *Rickettsia* genus. The exception was the *Francisella* genus, where almost all samples carried at least one ARG (98.9%). The resistance genes predicted for each sample are shown in Supplementary File 2. Detected ARGs for each species are summarized in Table 2.

Although there is a high number of ARG-positive samples from the *Francisella* genus, a large part of them (791 samples) was predicted to contain only one ARG. Furthermore, all of these samples had the *FTU-1* gene as the one predicted resistance gene. *FTU-1* was very prevalent in the analysed *Francisella* samples as all positive ones harboured this gene (948 samples out of the total 959). This gene is a class A *β* -lactamase gene found in almost every *Francisella tularensis* genomes^32^, including the reference genome of the species (based on the analysis with the same pipeline of the *F. tularensis* reference genome assembly available at NCBI: https://www.ncbi.nlm.nih.gov/assembly/GCF_000008985.1/). As a consequence of the high prevalence of this gene, we have additionally calculated the ratio for the *Francisella* samples containing ARGs without taking *FTU-1* gene into consideration. The percentage of positive samples without this gene is 16.3%, which is still the highest ratio found in our analysis. The number of ARG positive and negative samples can be found in Table 1. Supplementary File 2 shows all the resistance genes predicted by RGI for each positive biosample included in our analysis.

**Table 1.**
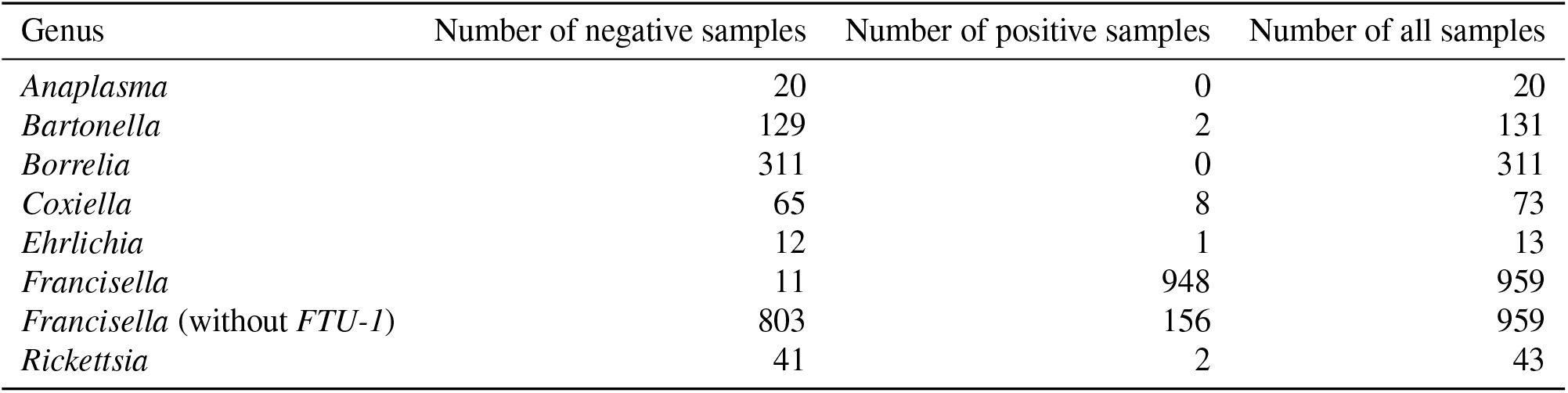
ARG positive and negative samples by the genus of origin.

| Genus | Number of negative samples | Number of positive samples | Number of all samples |
| --- | --- | --- | --- |
| *Anaplasma* | 20 | 0 | 20 |
| *Bartonella* | 129 | 2 | 131 |
| *Borrelia* | 311 | 0 | 311 |
| *Coxiella* | 65 | 8 | 73 |
| *Ehrlichia* | 12 | 1 | 13 |
| *Francisella* | 11 | 948 | 959 |
| *Francisella* (without *FTU-1*) | 803 | 156 | 959 |
| *Rickettsia* | 41 | 2 | 43 |

**Table 2.**
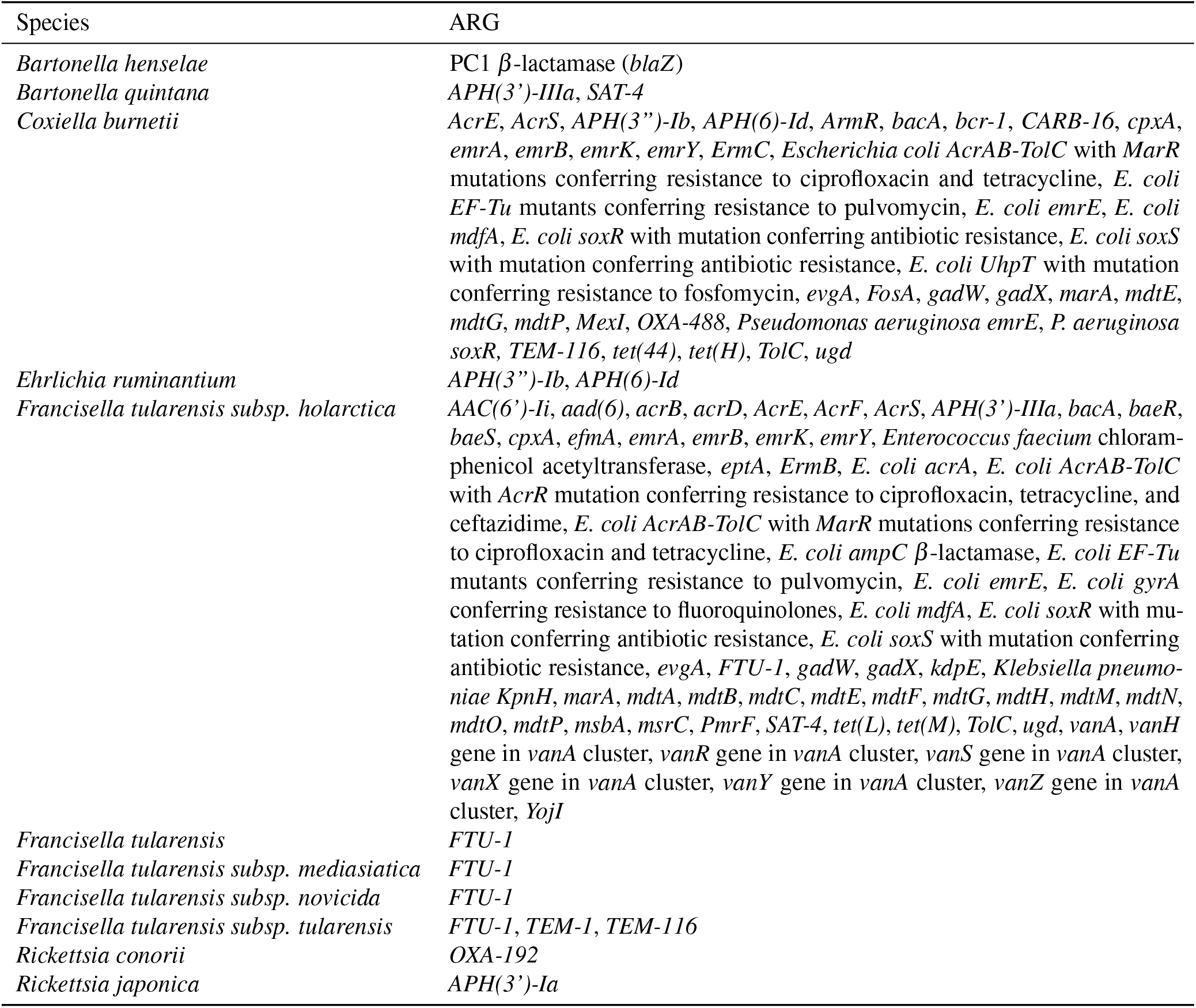
Resistance genes detected for each species by RGI and CARD. In case of *Francisella tularensis*, the samples were listed at the subspecies level, where that information was available, otherwise, they were summarized at the species level.

### Mobility analysis

Contigs, where ARGs were predicted by RGI, were further analysed to determine if these genes could be associated with any mobile genetic elements. An ARG was considered potentially mobile if the contig it was found on was predicted to be of plasmid origin, if it was within a predicted phage sequence or if it was found within 10098 bp distance from the closest detected iMGE (for details on the mobile element determination and the selection of the cut-off value see Materials and Methods). The number of potentially mobile and immobile ARGs can be found for each genus in Table 3. Further details of mobile elements associated with each predicted ARG are presented in Supplementary File 2.

**Table 3.**
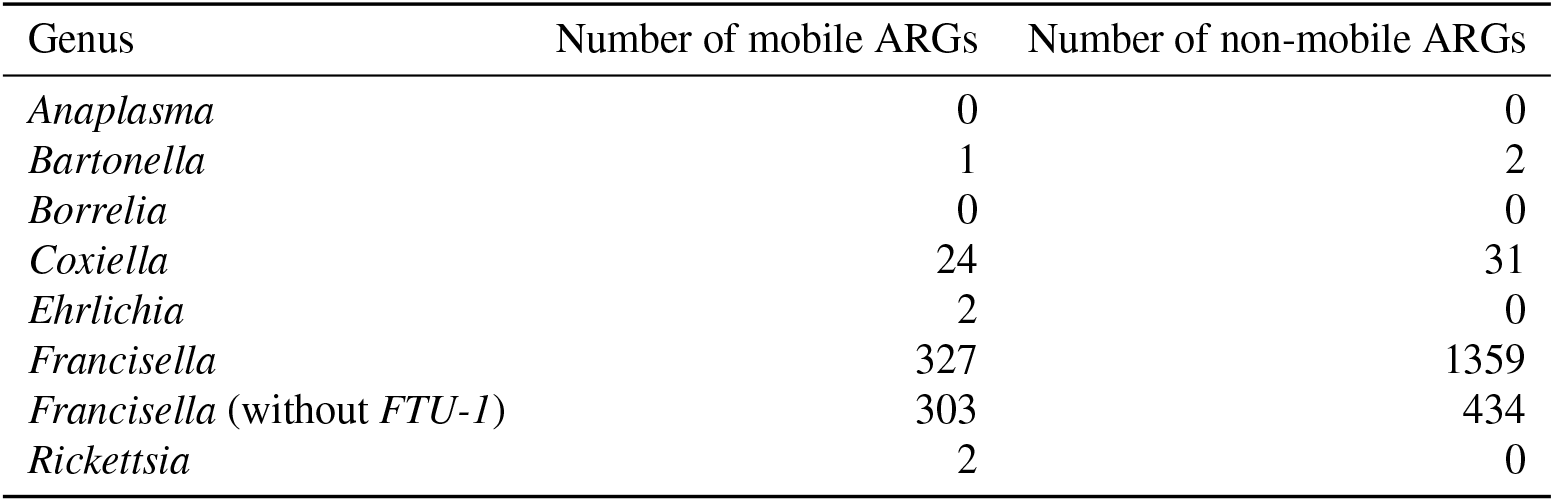
Number of mobile and immobile ARGs for each genus analysed.

### Comparing species and environment of origin between ARG positive and negative samples

Understanding the ecological environment where the samples included originated from is useful to gain a deeper insight into the potential factors underlying our results. The background of the samples, either positive or negative considering their ARG content, was assessed based on the origin of samples and species of the isolates. Samples were enrolled into origin categories based on their metadata (for details, see Materials and Methods). The ARG positive and negative sample distribution among the categories of origin are summarized in Table 4.

**Table 4.**
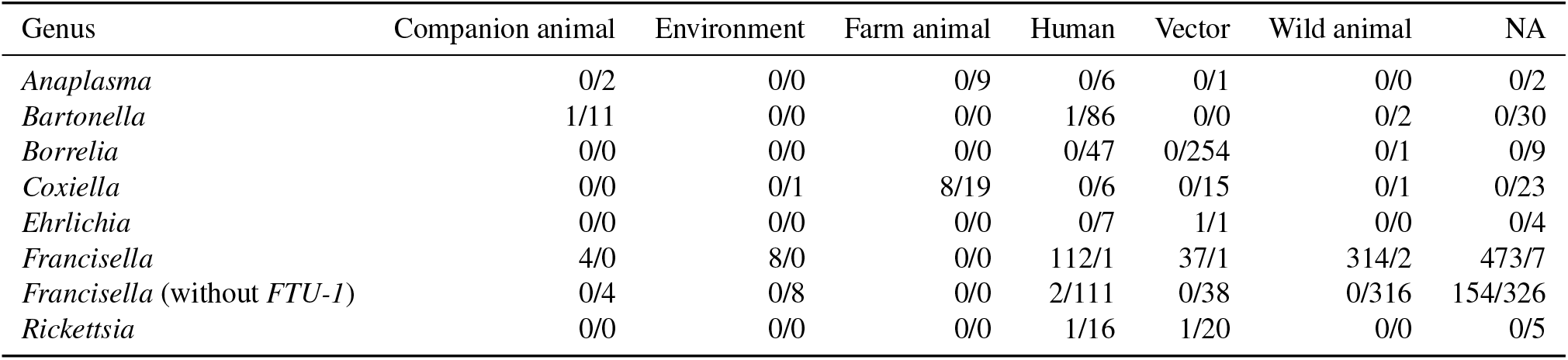
Number of ARG positive/negative samples in different origin categories.

Interestingly, only samples originating from farm animals were positive to ARGs in the case of *Coxiella burnetii*, while such high dependence wasn’t revealed for any other genus. To compare the association of positive samples to farm animal origins at *Coxiella* genus and all the remaining samples (excluding *Francisella*), Fisher’s exact tests were performed on the contingency tables from Figure 2. A significant difference was revealed in the case of *Coxiella* (OR: Inf, 95% CI: 3.6-Inf, p<0.001), however, such differences could not have been found in case of the rest of the samples (OR: Inf, 95% CI: 0.01 - Inf, p > 0.9999).

**Figure 2.**
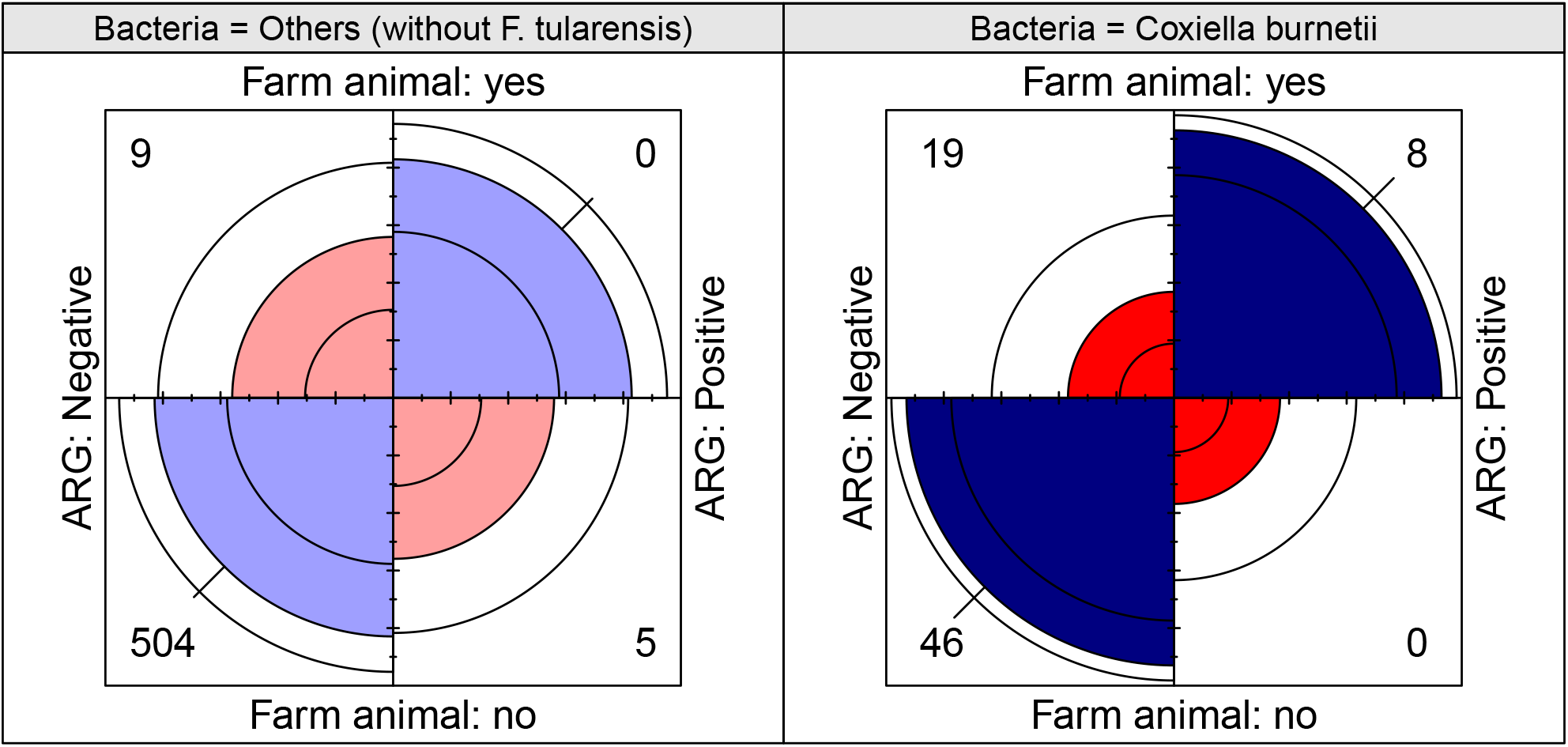
Fourfold displays showing the difference between ARG positivity and farm animal origin, in the case of *Coxiella burnetii* (right-hand side) and the rest of the samples (without *Francisella* genus, left-hand side). A significant imbalance (p<0.001) is apparent in the case of *Coxiella*, as the Fisher’s tests showed (see the Results section for details).

The species of origin of ARG positive and negative samples are also of a key importance to understand the presence of genotypic resistance among microorganisms from different ecological environments. The number of ARG positive and ARG negative samples from each species can be found in Supplementary File 3.

### Comparison between different methods

To understand the effect of the applied methods on our observations, we have complemented the analysis by including two additional methods. A similar approach to RGI is the NCBIs AMRFinderPlus pipeline which uses the NDARO database for ARG annotation. This tool utilizes a separate (the so-called plus database) for virulence-, metal- and biocide resistance genes. Even though CARD stores information regarding disinfectants, we have not used the plus genes in our analysis with AMRFinderPlus to keep as much consistency between the databases as possible. Furthermore, to compare the results from a pipeline with a fundamentally different approach, KmerResistance was also employed on the raw data with both the CARD and NDARO databases. For consistency to the analysis with AMRFinderPlus, plus genes were removed from the results of the KmerResistance analysis with the NDARO database. (Further details on the analysis pipelines are presented in the Materials and Methods section.)

Due to the inherent differences between databases and methods, their results are not directly comparable in a study like ours. However, to still augment the confidence of our findings, we compared the ARG positive samples among the four methods. Venn diagrams of ARG-positive samples from the so-called methodologies for each genus incorporated in the analysis can be found in Supplementary Figures 1-8 (Supplementary File 4). Detailed ARG predictions from each method can be found in Supplementary Files 5-7. Results for *Anaplasma, Borrelia, Coxiella, Ehrlichia* and *Rickettsia* genera were in line by each employed method, with differences affecting only a few samples (Supplementary Figures 1, 3-5 and 8 in Supplementary File 4). In the case of the genus *Bartonella*, KmerResistance with CARD and NDARO databases has predicted an additional 13 and 11 ARG positive samples compared to RGI, respectively (Supplementary Figure 2 in Supplementary File 4). The short read-based approach predicted more positive samples in the case *Francisella* genus as well (when FTU-1 gene wasn’t considered), where KmerResistance with CARD found 60 and KmerResistance with NDARO identified 34 ARG-positives more (Supplementary Figure 7 in Supplementary File 4).

## Discussion

We have revealed a lack of ARGs in the available and high-quality WGS samples of human-pathogenic and potentially tick-borne bacteria. *Francisella* showed the highest ratio of samples with at least one ARG (16.3%), however, ARG-positivity was generally very low in the rest of the analysed genera (only 2.2 %). This finding is surprising at first, considering the vast amount of antibiotic pollution and ARG prevalence of the natural and artificial environments^3–7,33–35^. The effect of natural environments on the spread of antimicrobial resistance (AMR) is well represented by the fact that wild animals were found to harbour different ARGs^36,37^, even though it might correlate with anthropogenic influence^38,39^. Furthermore, it is well known that antibiotic resistance is not only dependent on human activity as its origins root long before the onset of clinical antibiotic usage^40–42^. Considering this widespread occurrence of resistance, we believe that the potential factors influencing our findings deserve a closer look.

Our analysis has included data generated by second and third-generation sequencing technologies. Even though these methods have proven useful in numerous research areas, including antimicrobial resistance, they also have certain limitations. For example, if some genomic elements (e.g. ARGs) are absent from the sample, as in our case, only with sufficient sequencing depth can one conclude that true biological variation is the reason for the missingness^43^. To prevent such artefacts in our results, we have limited our analysis to samples with at least 95% coverage to their respective representative genome. Furthermore, we have used a 90% coverage and identity threshold for filtering the raw ARG predictions for limiting potential false positive calls. Another potential source of bias in genome-based resistance gene surveillance studies is that the results can only depend on the currently available resistance gene resources and tools, that might have many differences among them^44^. To mitigate these effects on our results, the samples included in our analysis were analysed with two additional tools and one additional database. NCBI’s AMRFinderPlus tool with the NDARO database and the short read-based KmerResistance tool with both the CARD and NDARO databases were employed, besides the RGI with CARD database approach. These results indicate that RGI was highly concordant with the additional methods used for the analysis, even though KmerResistance has predicted more positive samples for the *Bartonella* and *Francisella* genera. It can be assumed that short-read technologies are more sensitive, which could explain why more positive samples were found with KmerResistance; however, these methods have not been systematically compared yet.

Phenotypic resistance to various antibiotics has been described in the bacteria we have studied, but to the best of our knowledge, multi-drug resistant strains have not yet been found. Interestingly, the first choice for treatment of infections caused by the the majority of bacteria included in our study is doxycycline^13,45–53^. For *F. tularensis* aminoglycosides and fluoroquinolones are also commonly recommended^54^. Resistance to fluoroquinolone antibiotics has been identified in *Bartonella, Borrelia* and *Ehrlichia* species^47,52,53^. Nevertheless, fluoroquinolone resistance in *Ehrlichia* spp. is probably due to intrinsic effects^55^. In *Ehrlichia*, resistance to macrolides, aminoglycosides and certain *β* -lactams has also been detected^47,56^, which may be consistent with the aminoglycoside resistance genes we found in *E. ruminantium* (Table 2). In contrast to the lack of ARGs in *Borrelia* species involved in our study, *β* -lactam and rifampicin resistance have already been described for them^51,52^ (Table 2). However, it should be noted that using the two KmerResistance-based approaches, we did find an aminoglycoside resistance gene and an efflux pump in this group (*APH(3’)-Ia, E. coli emrE*, Supplementary Files 6-7). *C. burnetii* and *F. tularensis* samples were found to encode a wide range of resistance genes in our analysis (Table 2). While phenotypically manifested ciprofloxacin and chloramphenicol resistance have already been described in *C. burnetii*^49^, furthermore, resistance to penicillin has also been described^57^, several strains appear to show susceptibility to ampicillin^49^. Moreover, in *F. tularensis*, resistance to *β* -lactam, macrolides, linezolid and clindamycin was described previously^58–60^.

However, it is important to note that the resistance gene content of the genome does not necessarily correlate with phenotypic resistance^61^, and in any case, our results will be limited by the capacity of databases and tools^44,62–64^. Furthermore, there are many factors affecting the in vivo susceptibility of a microorganism, compared to what was discovered in vitro^53,65^. In addition, it should be noted that of the 156 ARG positive *F. tularensis* samples in our analysis (when the *FTU-1* gene was excluded), 153 were from BioProject PRJNA669398. In this BioProject, samples derived from the live vaccine strain of *F. tularensis subsp. holartica* (LVS) and were tested for the presence of evolutionary mutations under the pressure of doxycycline and ciprofloxacin antibiotics^66^. However, it is interesting to note that of the strains included in our study and identified as LVS based on the available metadata, 43.7% (153/350) of the strains belonging to the BioProject PRJNA669398 were found to be ARG positive (without *FTU-1*), while 0% (0/49) of the LVS strains included in other BioProjects were positive. These aspects raise the question of the reliability of the samples from this BioProject.

The general consensus is that de novo resistance develops in response to selective pressure from antibiotics or potentially other stressors (e.g. heavy metals), which is then associated with resistance mutations or the uptake of acquired ARGs via horizontal gene transfer^67–69^. Consequently, it is important to consider the ecological environment of microorganisms to investigate the underlying effects of the resistance gene deficiency we have observed. Most microorganisms we studied are obligate intracellular bacteria, strictly bound to vectors and hosts^46,53,70–76^. Exceptions are *Borrelia* species, which despite being strongly associated with hosts- and vectors are not intracellular pathogens^77^, and the *Francisella* and *Coxiella* genera, where transmission pathways outside of vectors are also important^78,79^. Consequently, there is a limited medium in which selective pressure and the presence of ARGs would co-occur. A tick or possibly another arthropod vector could theoretically be subjected to more or less antibiotic exposure in its environment, for example, when it takes up fluid from plants or soil, but the extent to which this effect could represent a true selective force is questionable. In addition, the presence of available ARGs might also be limited, as it has been described that the ticks’ own bacteriota limits the colonisation of microorganisms from the environment^80^.

Bacteria in hosts can be subjected to more pronounced antibiotic pressure. Of course, this is not primarily in wild animals, but when bacteria are introduced into domestic animals or even humans. It is assumed that a significant proportion of human infections are treated with antibiotics, and even if this were to involve the uptake of ARGs of these strains, it is unlikely that the bacteria that have acquired resistance genes would be re-introduced into a tick. Domestic animals may be of much greater importance in this respect. In their case, antibiotic treatment is also significant, or perhaps even more significant, while a vector may also go undetected more easily. Farm animals may be particularly noteworthy in this respect. However, it is interesting that we found a strong concordance between ARG positivity and the farm animal origin of the samples only in the case of *C. burnetii* (Figure 2). One could hypothesize that the majority of *C. burnetii* samples from farm animals were from intensive keeping conditions, whereas the opposite was true for samples from other genera. Cattle can be considered as the classic intensively kept animal in contrast to goats, sheep and horses. For *C. burnetii* almost half of the 27 samples from farm animals were derived from cattle (14), of which 5 were positive, and there were 3 additional positive samples from goats (Supplementary Files 1, 2). It is important to note, however, that one positive sample in the genus *Ehrlichia* was from a tick collected directly from a sheep (BioSample: SAMN04335506, Supplementary File 1). Nevertheless, this was classified of vector origin in our analysis as it is not known if the tick was infected a priori. It should also be noted that the *Anaplasma* genus also included 9 samples of farm animal origin, 6 of which were of cattle origin, 1 of equine origin and 2 of sheep origin, but none of these was found to be positive (Supplementary Files 1, 2). Our reasoning is, of course, limited by the availability of metadata, but it does not appear that the confounding effect of cattle, as traditionally intensively kept animals, is the cause of the phenomenon observed in the case of *Coxiella*. A difference between the *Anaplasma* and *Coxiella* genera may be due to their life cycle. Namely, the extracellular form of *Coxiella* also plays an important role in its spread, and consequently, it is not entirely bounded to vectors^78^. Of course, the ability of the environmental form of *C. burnetii* to take up genes from the environment and the extent to which these might be necessary for its survival in the presence of antibiotics is questionable.

*Francisella tularensis* is similar to *C. burnetii* in some respects. The role of vectors in its spread is not necessarily exclusive^79^, and it can occur in the infected hosts in an extracellular state^81,82^. Several ARG positive samples of *F. tularensis* were found, but these were almost exclusively from an experiment in which artificial antibiotic pressure was applied to the bacteria, making these results difficult to interpret (see above). Furthermore, it is, of course, possible that the microorganisms studied do acquire resistance genes. However, these were impossible to detect because they were dropped by the bacteria. This may be because it is assumed that the tick’s gut provides a fundamentally different microenvironment for the bacteria than the circulation of the host animal, including the existing selection pressure. The continuous activation of resistance mechanisms is an energy-demanding process, so without therapeutic pressure, these costs exceed the benefits. Thus, the bacteria try to get rid of them, which may explain the lack of ARGs. This effect has been described for ARGs; however, the fitness cost might depend on the resistance mechanism^67,83,84^.

Our results show that the potentially tick-borne and human pathogenic microorganism strains we investigated lack antimicrobial resistance genes. Comprehending the processes underlying this phenomenon may be an important aspect in understanding the ecology of these species and, through this, in assessing the risk of phenotypic antibiotic resistance in clinical settings. However, an exception to the above statement of the tick-borne human pathogens is *Coxiella burnetii*, where resistance is a potential veterinary medical and public health threat when farm animals and their products are considered.

## Supplementary information

**Supplementary File 1**: Tables for the metadata on each samples analysed in this study. Samples from the included genera are placed in separate sheets in the excel file. The tables contain the NCBI BioSample ID (column BioSample ID), the sample origin (column sample_origin), sample host (column sample_host), country (column country) and species (column species) metadata information available for each sample at the NCBI SRA database, the origin category (column Origin) samples were enrolled based on the available metadata and if the sample was included in the analysis with the KmerResistance tool (column kmerresistance).

**Supplementary File 2**: Tables for the predicted ARGs by RGI. Results for each genera are located in separate sheets in the excel file. The columns correspond to the BioSample ID (column BioSample ID), the ARGs name as in the CARD database (column Best_Hit_ARO), the identity and coverage estimated by RGI (columns Identity and Coverage), the gene family available for the ARG in the CRAD database (column AMR Gene Family), the predicted plasmid origin of the contig the ARG is located at (column Contig Plasmid Origin), the type of the phage sequence if the ARG was found within one (column Associated Bacteriophage Sequence) and the integrative mobile genetic elemnt the ARG was found to be associated with (column iMGE).

**Supplementary File 3**: Tables for the number of positive and negative samples for each species. Separate sheets are used for each genera included in our study. In the case of *Francisella tularensis* each supspecies was considered separately if that data was available. Positive and negative samples were counted for *F. tularensis* with and without considering FTU-1 as well. Columns correspond to the name of the species (column Species), the number of negative samples (column Number of negative samples) and the number of positive samples (Number of positive samples).

**Supplementary File 4**: Supplementary Figures 1-8 and their associated legends.

**Supplementary File 5**: Tables for the results of the AMRFinderPlus tool. Results for each genera are located in separate sheets in the excel file. The columns correspond to the BioSample ID (column BioSample ID), the gene symbol and sequence name as availabel in the NDARO database (columns Gene symbol and Sequence name) and the coverga and identity as estimated by the AMRFinderPlus tool (columns Coverage and Identity).

**Supplementary File 6**: Tables for the results of the KmerResistance tool when used with the CARD database. Results for each genera are located in separate sheets in the excel file. The columns correspond to the BioSample ID (column BioSample ID), the header for the ARG in the CARD protein homolog model database fasta file (column CARD database fasta header) and the template length, template coverage, template identity, query identity and query coverage as presented by the KmerResistance tool (columns Template_length, Template_Identity, Template_Coverage, Query_Identity and Query_Coverage).

**Supplementary File 7**: Tables for the results of the KmerResistance tool when used with the NDARO database. Results for each genera are located in separate sheets in the excel file. The columns correspond to the BioSample ID (column BioSample ID), the header for the ARG in the NDARO database fasta file (column NDARO database fasta header) and the template length, template coverage, template identity, query identity and query coverage as presented by the KmerResistance tool (columns Template_length, Template_Identity, Template_Coverage, Query_Identity and Query_Coverage).

**Supplementary File 8**: Tables for the NCBI BioProject and corresponding BioSample IDs of the samples that were included in the present analysis. Samples from the included genera are placed in separate sheets in the excel file. The columns correspond to the NCBI BioProject ID (column BioProject ID) and the comma separated list of the associated NCBI BioSample IDs (column BioSample ID) that were included in our analysis.

## Declarations

### Ethics approval and consent to participate

Not applicable.

### Consent for publication

Not applicable.

### Availability of data and material

Datasets analysed in the present study are available at the National Center for Biotechnology Information (NCBI) Sequence Read Archive (SRA). For more information on the BioSample IDs and the corresponding BioProject of each analysed samples, please see the Supplementary File 8.

### Author contributions statement

NS takes responsibility for the data’s integrity and the data analysis’s accuracy. MP and NS conceived the concept of the study. MP, NS and SÁN participated in the bioinformatic analysis. MP and NS participated in the drafting of the manuscript. AGT, GV, LM, MP, NS, RF and SÁN carried out the critical revision of the manuscript for important intellectual content. All authors read and approved the final manuscript.

## Acknowledgements

The authors would like to thank the providers of each BioProject employed in the present study.

## Authors’ information

No relevant information can be provided of the authors that may aid the readers’ interpretation of the article.

