## Supplementary material for "Antimicrobial resistance gene lack in tick-borne pathogenic bacteria": SupplementaryFile_4.pdf

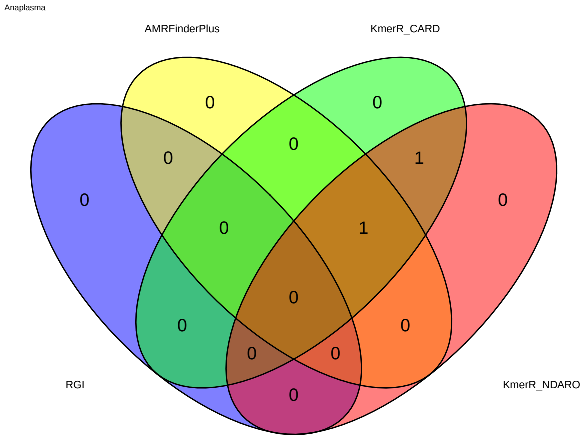


**Supplementary Figure 1: Venn diagram of ARG positive samples between the four methods for Anaplasma genus.** The agreement between the RGI, AMRFinderPlus and KmerResistance with CARD (KmerR_CARD) and NDARO (KmerR_NDARO) databases is represented as a Venn diagram. A sample was considered to be positive if it was predicted to have at least one ARG by the tools, respectively. For details on the four methods, please refer to the Materials and Methods section.

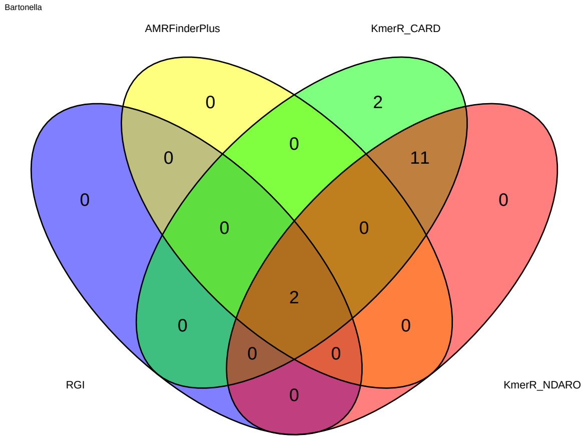


**Supplementary Figure 2: Venn diagram of ARG positive samples between the four methods for Bartonella genus.** The agreement between the RGI, AMRFinderPlus and KmerResistance with CARD (KmerR_CARD) and NDARO (KmerR_NDARO) databases is represented as a Venn diagram. A sample was considered to be positive if it was predicted to have at least one ARG by the tools, respectively. For details on the four methods, please refer to the Materials and Methods section.

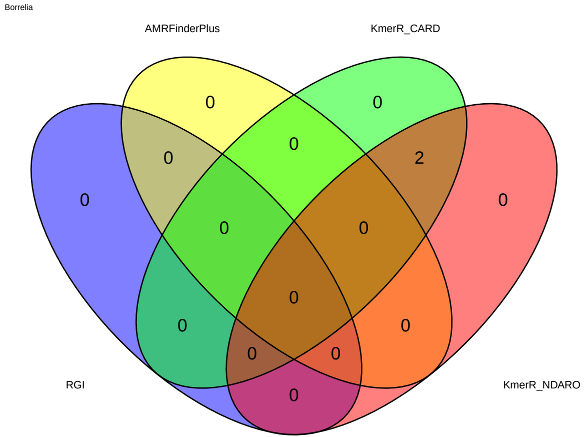


**Supplementary Figure 3: Venn diagram of ARG positive samples between the four methods for Borrelia genus.** The agreement between the RGI, AMRFinderPlus and KmerResistance with CARD (KmerR_CARD) and NDARO (KmerR_NDARO) databases is represented as a Venn diagram. A sample was considered to be positive if it was predicted to have at least one ARG by the tools, respectively. For details on the four methods, please refer to the Materials and Methods section.

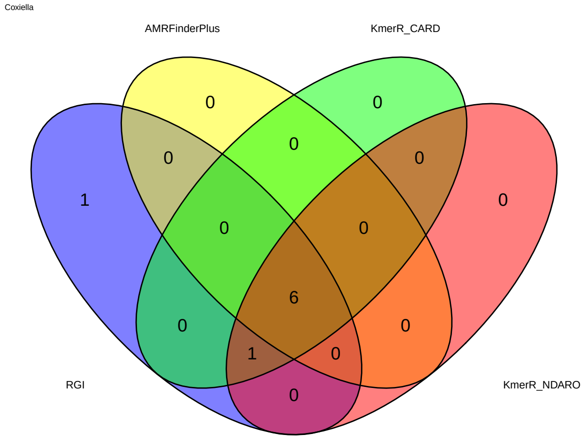


**Supplementary Figure 4: Venn diagram of ARG positive samples between the four methods for Coxiella genus.** The agreement between the RGI, AMRFinderPlus and KmerResistance with CARD (KmerR_CARD) and NDARO (KmerR_NDARO) databases is represented as a Venn diagram. A sample was considered to be positive if it was predicted to have at least one ARG by the tools, respectively. For details on the four methods, please refer to the Materials and Methods section.

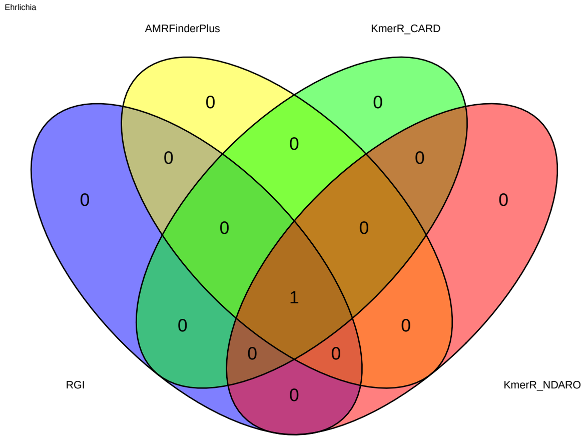


**Supplementary Figure 5: Venn diagram of ARG positive samples between the four methods for Ehrlichia genus.** The agreement between the RGI, AMRFinderPlus and KmerResistance with CARD (KmerR_CARD) and NDARO (KmerR_NDARO) databases is represented as a Venn diagram. A sample was considered to be positive if it was predicted to have at least one ARG by the tools, respectively. For details on the four methods, please refer to the Materials and Methods section.

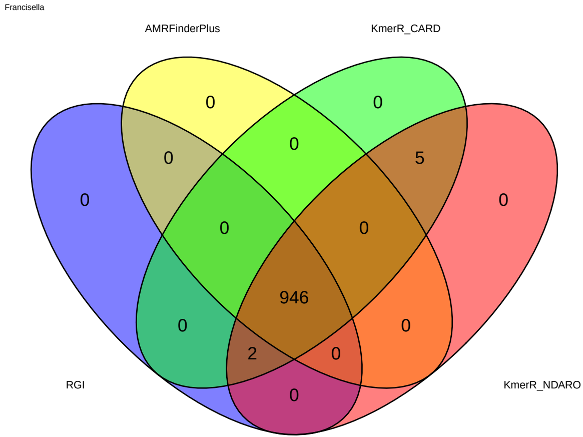


**Supplementary Figure 6: Venn diagram of ARG positive samples between the four methods for Francisella genus.** The agreement between the RGI, AMRFinderPlus and KmerResistance with CARD (KmerR_CARD) and NDARO (KmerR_NDARO) databases is represented as a Venn diagram. A sample was considered to be positive if it was predicted to have at least one ARG by the tools, respectively. For details on the four methods, please refer to the Materials and Methods section.

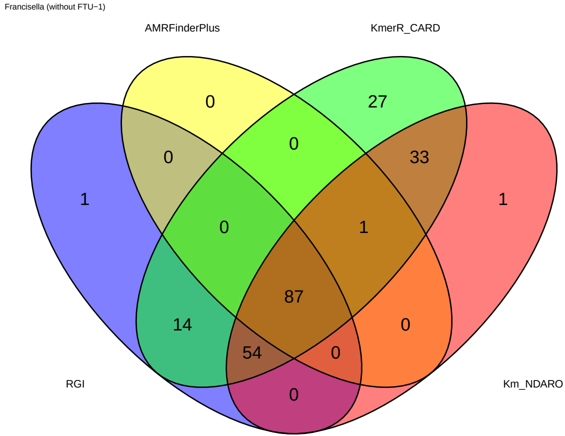


**Supplementary Figure 7: Venn diagram of ARG positive samples between the four methods for Francisella genus (without the FTU-1 gene).** The agreement between the RGI, AMRFinderPlus and KmerResistance with CARD (KmerR_CARD) and NDARO (KmerR_NDARO) databases is represented as a Venn diagram. A sample was considered to be positive if it was predicted to have at least one ARG by the tools, respectively. For details on the four methods, please refer to the Materials and Methods section.

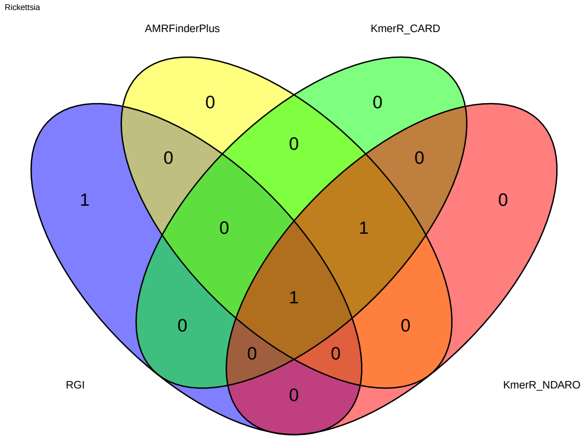


**Supplementary Figure 8: Venn diagram of ARG positive samples between the four methods for Rickettsia genus.** The agreement between the RGI, AMRFinderPlus and KmerResistance with CARD (KmerR_CARD) and NDARO (KmerR_NDARO) databases is represented as a Venn diagram. A sample was considered to be positive if it was predicted to have at least one ARG by the tools, respectively. For details on the four methods, please refer to the Materials and Methods section.
